# Involuntary facial muscle activity during imagined vocalisation contaminates EEG and enables emotion decoding

**DOI:** 10.64898/2026.03.18.712559

**Authors:** Yichen Tang, Paul M. Corballis, Luke E. Hallum

## Abstract

**Objective:** Decoding imagined speech from electroencephalography (EEG) recordings is potentially useful for brain-computer interfaces. Previous studies have focused on decoding semantic information from EEG, leaving the decoding of emotion – an important component of human communication – largely unexplored. *Approach*. Here, we report two experiments involving participants tasked with overt (n = 14) or imagined (n = 21) emotional vocalisation in five different categories: anger, happiness, neutral, sadness, and pleasure. Throughout, we recorded 64-channel EEG; we computed time-frequency features and used a logistic-regression classifier to evaluate emotion decoding accuracy. In five participants, we also recorded facial surface electromyography (sEMG) during imagined vocalisation, and studied the contamination of EEG by sEMG. *Main results*. Our results show that emotion can be decoded from single-trial EEG recordings of both overt (78.1%, chance = 20%) and imagined vocalisation (36.4%). The high-gamma band (50 to 100 Hz) and lateral EEG channels (T7, T8, and proximal) were important for decoding. sEMG analysis indicated that involuntary facial muscle activity contributed to these spectral and spatial patterns during imagined vocalisation. We identified a prominent muscle-induced pattern over lateral EEG channels – the railroad cross-tie pattern – showing greater activation during imagined happy vocalisations. We conclude that involuntary facial muscle activity is associated with certain emotion categories (i.e., happiness) and contributes to the decoding of emotion from single-trial EEG recordings. *Significance*. This study provides converging EEG and sEMG evidence demonstrating that involuntary facial muscle activity occurs during imagined vocalisation and is modulated by emotion, producing emotion-specific EEG contamination patterns. Although we focused on imagined vocalisations, the EMG patterns we observed closely resembled EEG patterns commonly reported in emotion decoding studies, raising concerns about involuntary facial muscle activity in such work.

## 1 Introduction

A major goal of research in brain computer interfaces (BCIs) is to build direct communication pathways between the human brain and external devices, allowing humans to control machines without relying on voluntary muscle movements [1]. Among various techniques for detecting brain activity in BCI systems, electroencephalography (EEG) is most commonly used due to its high coverage of cortical regions, safety, accessibility, portability, cost-effectiveness, and high temporal resolution [2, 3]. However, there are many significant challenges for EEG-based BCI systems to establish an effective communication pathway. One major challenge is undesired physiological contaminations reflected in EEG, such as facial and neck muscle activity, blinks and eye movements, skin potentials, and carotid pulse, resulting in degraded BCI performance and unreliable analyses [4, 5, 6]. Among these artefacts, muscle activity (electromyography, EMG) is particularly difficult to separate from EEG due to its wide spatial spread over the scalp and its spectral overlap with neural EEG signals [6, 7]. In EEG-based BCI studies, possible EMG contamination has often been ignored [8]. This can, of course, lead to misinterpretations in which the EMG artefact is misattributed as a brain signal. Here, we explore the influence of EMG on decoding emotion during imagined speech.

Humans infer not only semantic information from speech, but also paralinguistic features such as identity, age, gender, and the speaker’s current emotional state [9]. In the past decade, EEG has been used to decode semantic content from imagined speech, aiming at binary or multi-class classifications of a selected set of vowels [10, 11], syllables [12, 13], or words [11, 13, 14, 15, 16, 17]. Progress has been made, with multiple studies showing accuracies *>*80% decoding binary classes [11, 10, 17] and accuracies significantly above chances for multi-class classifications (e.g. 61.4% for classification of five words, chance=20% [16]; 57.1% for classification of seven syllables and four words, chance=9% [13]). However, despite this progress, an equally important aspect in human communication – emotion – has received comparatively little attention in imagined speech EEG studies [18, 19]. Decoding emotion from imagined speech could provide BCI systems additional information to make affect-aware, more user-friendly decisions, and to enhance human-to-human communications through these systems.

There have been EEG-based studies on the overt production of emotional speech [20, 21, 22]. Rohr et al. tasked participants with pronouncing emotional words following a visual cue, and observed differences in event-related potential (ERP) amplitudes between negative and neutral or positive emotional words [20]. Both Abtahi et al.[21] and Ghoniem et al. [22] used machine learning methods for decoding emotion from overt speech. Abtahi et al. tasked participants with pronouncing an emotionally neutral sentence “The sky is green” with seven different tones, corresponding to seven emotions (happiness, sadness, anger, surprise, fear, disgust, and neutral) [21]. The authors extracted time-frequency features from EEG recorded during speech production and applied a neural network to decode emotion across participants, resulting in a decoding accuracy of 66.4% (chance=14%) [21]. Ghoniem et al. tasked participants with discriminating song clips into one of seven emotion categories (fear, surprise, happiness, disgust, neutral, anxiety, and sadness) and vocally describing their feelings (e.g., “I am feeling deep depression”) while listening to the clips [22]. The authors extracted a large set of temporal, spectral, and time-frequency EEG features and applied a novel neural network model to decode emotion, reaching an exceptionally high emotion classification accuracy of 98.06% (chance=14%) across participants [22].

These studies took different approaches for removing EMG contamination from facial and articulatory muscle activity during overt speech. Abtahi et al. [21] and Ghoniem et al. [22] simply applied band-pass filters at 0.1-30 Hz and 0.5-64 Hz, respectively. However, due to the wide spectrum covered by EMG activity [7, 23, 24, 25, 26], band-pass filters cannot remove all EMG contamination, and EMG activity is likely to have biased their decoding performance. Rohr et al. employed a residue iteration decomposition (RIDE) algorithm [27, 20]. The method unmixes EEG signals into components synchronised to vocal onsets and those synchronised to stimulus onsets (i.e. cue for starting pronunciation). It then rejects vocal onset-synchronised components, which are assumed to be articulation artefacts dominated by EMG, and remixes the signals [27]. This method is claimed to have effectively isolated the speech artefact, but its usage is limited to the analysis of stimulus-evoked, synchronised ERP responses, and is not applicable for online single-trial EEG decoding.

Recent studies have begun to examine emotion-related EEG features during imagined speech. Kapitány-Fövény et al. [18] differentiated participants with depression from controls using EEG recorded during both imagined and overt production of emotion words (positive, neutral, negative). Roh et al. [19] performed binary valence classification (positive versus negative) from EEG recorded during imagined production of emotional sentences and analysed the contribution of different scalp regions to decoding performance. Neither study addressed the possibility of EMG contamination, reflecting the common assumption that imagined speech, which involves no voluntary facial or articulatory movement, is relatively free of myogenic artefacts. This assumption is increasingly challenged by converging surface electromyography (sEMG) evidence from imagined speech and related states. Nalborczyk et al. [28] reported elevated facial sEMG during induced verbal rumination; Bhadra et al. [29] showed that facial sEMG recorded during the production of imagined syllables contained decodable phonetic information. Recent imagined-speech EEG datasets have also begun to record facial sEMG together with EEG to enable EMG analyses [30, 31]. We hypothesise that involuntary facial muscle activity is also present during imagined emotional speech and contaminates EEG, and is modulated by emotion category.

In this study, we recorded multi-channel EEG during two experiments with participants tasked with overt and imagined production of emotional vocalisations. We performed within-participant emotion classification and examined the frequency bands and channels that enabled above-chance classifier performance. During the imagined vocalisation experiment, we also recorded facial surface electromyography (sEMG) from five participants in addition to the EEG channels and studied the correlation between facial muscle sEMG activity and EEG activity. We compared the frequency band and channel patterns for classification with the correlation between sEMG and EEG activity, to show whether muscle activity could result in the observed patterns. In addition, we performed emotion classification using a combination of EEG and sEMG features and compared the results to classification using only sEMG features. If a combination of EEG and sEMG features resulted in improved classification, this would suggest that EMG contamination had dominated EEG recordings and contributed to emotion decoding. Finally, we systematically detected and analysed one known pattern of EMG contamination – the “railroad cross-tie pattern” [7] – which we hypothesised to present during imagined vocalisation.

## 2 Methods

### 2.1 Experiments and data

#### 2.1.1 Overt and imagined vocalisation experiment design

We performed two experiments: one in which participants produced overt vocalisations (Experiment 1), and one in which they imagined vocalisations (Experiment 2). Fourteen participants (11 males and 3 females; aged 24 to 33 years, with an average age of 26.9 ± 2.75 years) and twenty-one participants (11 males and 10 females; aged 19 to 73 years, with an average age of 28.5 ± 12.82 years) with normal hearing and normal or corrected-to-normal vision participated in Experiments 1 and 2, respectively. Two older participants (59 and 73 years) increased the age range of the Experiment 2 cohort (19–32 years, mean 24.5 ± 3.7 after exclusion). Excluding these two outliers did not change the results of our analyses (Appendix A.1), so they were retained in all analyses. Each participant attended a single 3-hour session comprising six blocks, each of 50 trials. Participant P4 from Experiment 2 performed only five blocks of experiment due to a lack of time. This study was performed in accordance with the Declaration of Helsinki and approved by the University of Auckland Human Participants Ethics Committee (reference: UAHPEC24522). All participants provided written informed consent to participate in this study.

Figure 1 illustrates the trial structure. In Experiment 1, on each trial, the participant listened to an emotional vocalisation drawn from the Montreal Affective Voices (MAV) dataset [32] and recognised the emotion. Vocalisations were based on the vowel /ah/, and we included five emotion categories in this study: anger, happiness, neutral, pleasure, and sadness. After hearing the vocalisation, we provided the participant unlimited time to prepare their overt vocalisation of /ah/ using the recognised emotion. Once the participant was ready to vocalise, they pressed a key. We then provided a two-second countdown (during which the particpant viewed a central white circle, diameter = 0.1° of visual angle) to vocalisation onset. The vocalisation period lasted four seconds, however, the participants were instructed that they need not use the entire four seconds. Participants then provided a key press, indicating the recognised emotion (five-alternative forced-choice) and immediately thereafter received feedback against the labelled emotion in MAV. The participant proceeded from one trial to the next by a key press. To keep the participant engaged, we displayed their running score in the top left corner of the computer monitor (number of completed trials, number of correct trials).

**Figure 1.**
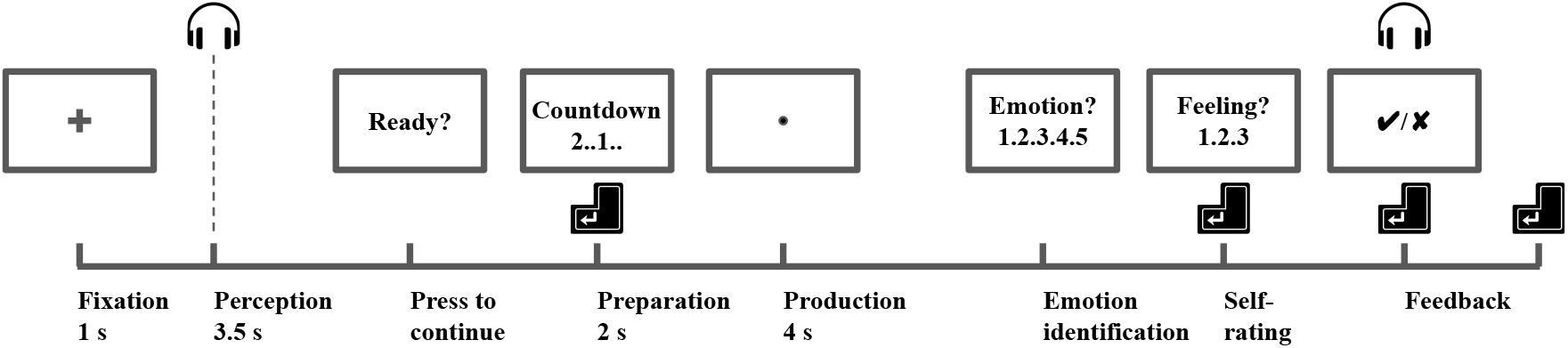
Trial structure for Experiment 1 and 2. The vocalisation stimulus was presented at the beginning of the “Perception” period; the participant was tasked with recognising the emotion (anger, happiness, neutral, pleasure, or sadness) and then, after a two-second countdown initiated by a key press, vocalising the recognised emotion (“Production” period). In Experiment 1, the participant overtly produced the vocalisation. In Experiment 2, the participant overtly produced the vocalisation on all trials in the first experimental block, and then, on all trials of the remaining five blocks, imagined producing the vocalisation. On each trial, the participant answered a five-alternative forced-choice question (“What was the recognised emotion?”) regarding the vocalisation stimulus (“Response”) and received feed-back, along with the correct answer (“Feedback”). In Experiment 2, additionally, the participant rated their feeling during the production of vocalisation (“Self-rating”).

The design of Experiment 2 was similar to Experiment 1. In Experiment 2, we tasked the participants to overtly produce vocalisations in the first block of trials, and to only imagine vocalisations in the remaining five blocks of trials. Specifically, we instructed the participants to imagine making the emotional vocalisations without reflecting the actions on external muscles (facial or articulatory). In Experiment 2, in addition to the five-alternative forced-choice question, we asked the participants to provide a subjective rating of their feeling during vocalisations, whether overt or imagined, to encourage emotion engagement. For example, if a participant perceived happiness in one trial, we asked the participant to choose a rating from: “very happy”, “moderately happy”, and “not happy”. All participants were instructed to keep their heads and bodies still, fixate on the centre of the screen, and try blink as little as possible during the experiment. We gave each participant 10 optional practice trials preceding each experiment block and several minutes of rest in between blocks.

#### 2.1.2 Visual and auditory setup

We performed the experiments in an electrically-shielded, quiet laboratory, dimly lit. We presented visual stimuli on a 24.5-inch OLED (organic light-emitting diode) display (PRM-224-3G-O; Plura Broadcast, Inc., Phoenix, Arizona, United States) with mean-luminance approximately 60 cd*/*m^2^, which we previously characterised using methods described in Hallum & Cloherty [33]. We delivered auditory stimuli through a pair of insert earphones (30+ dB external noise exclusion; ER-1; Etymotic Research, Inc., Elk Grove Village, Illinois, United States). The earphones were calibrated before each experimental session to produce a 1 kHz tone at 70±1 dB when coupled to an occluded-ear simulator (IEC 60318-4:2010 Ed.1.0 standard; calibrated at 1 kHz to match a KEMAR simulator, G.R.A.S. Sound & Vibration, ApS, Holte, Denmark).

#### 2.1.3 EEG, sEMG and speech recordings

We recorded 64-channel EEG using a BioSemi ActiveTwo AD-box (ADC-17; ActiveTwo; Biosemi, B.V., Amsterdam, Noord-Holland, Netherlands) at 2048 samples per second. We positioned the EEG electrodes according to the international 10-10 system (Figure 2-A). The participants’ vocalisations and recognised emotions were recorded along-side the EEG recordings. On five participants (P17 to P21) in Experiment 2, we also recorded four-channel bipolar sEMG using eight active Ag-AgCl electrodes, simultaneously with the EEG recordings. We placed the sEMG channels over multiple facial muscle groups, including the left medial frontalis, left temporalis, left orbicularis oculi, and left zygomatic major regions, as shown in Figure 2-B. For each bipolar electrode pair, an sEMG channel was derived by computing the voltage difference between the two electrodes. The activity of muscle groups close to the pair of bipolar electrodes was recorded by the sEMG channel. Under Experiment 1, we recorded the participants’ overt vocalisations using a dynamic microphone (SM58; Shure, Inc., Niles, Illinois, United States) placed approximately 10cm in front of the participant’s mouth; we time-locked this recording to EEG recordings through the Biosemi ActiveTwo AD-box.

**Figure 2.**
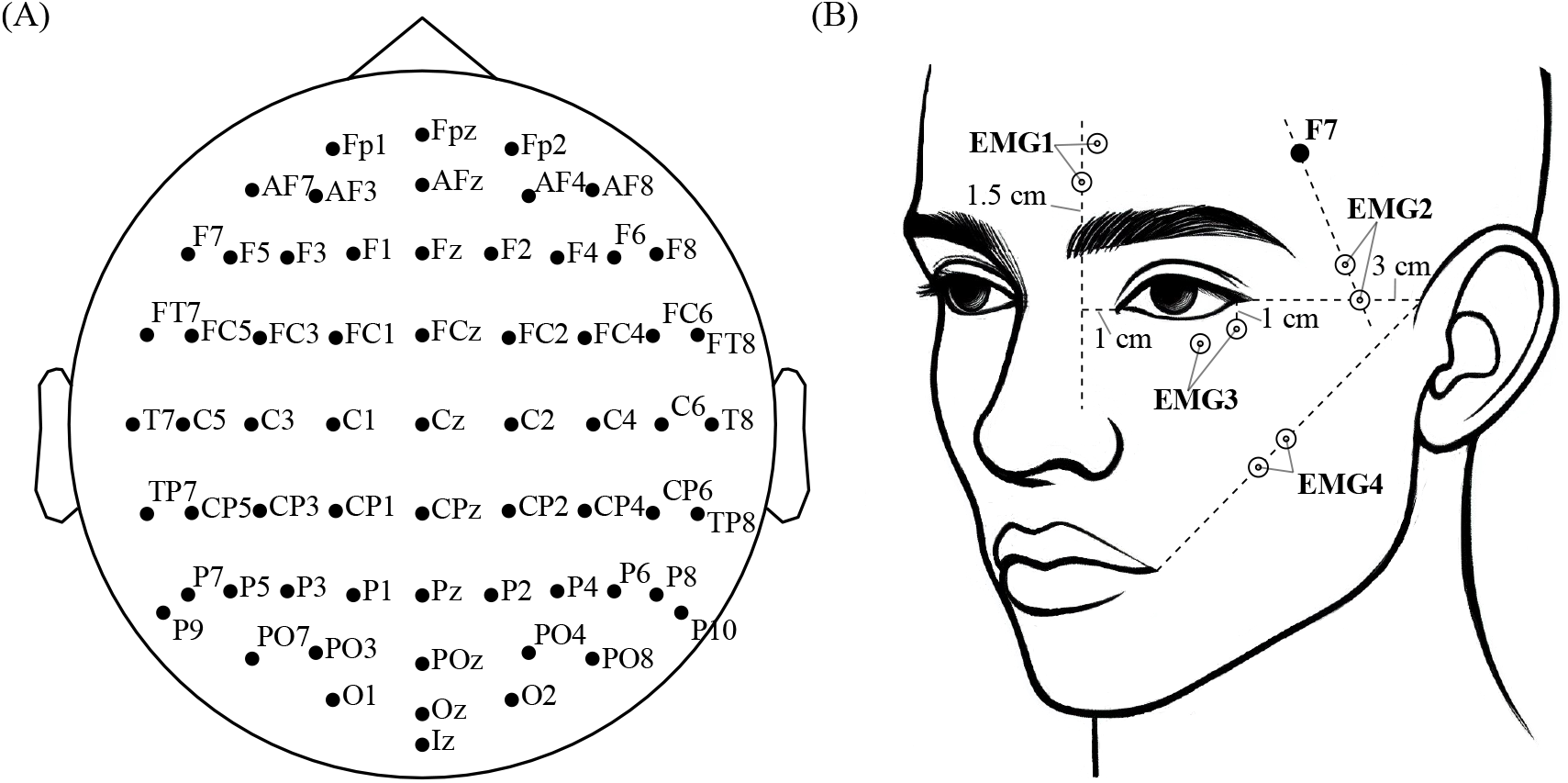
We recorded 64-channel EEG from all participants and, from five participants who performed imagined vocalisation, we simultaneously recorded EEG and four bipolar surface EMG (sEMG) channels. Panel (A) shows the EEG placement. Panel (B) shows the sEMG channel placement, following recommendations by previous studies [34, 7]. The distance between each pair of sEMG electrodes was set to 1.5 cm. EMG1 recorded muscle activity over the left medial frontalis region; EMG2 recorded activity over the left temporalis region; EMG3 recorded activity over the left orbicularis oculi region; and EMG4 recorded activity over the left zygomatic major region.

#### 2.1.4 Data preprocessing

We applied offline preprocessing steps using the open-source Python library MNE [35]. We applied a one-pass, zero-phase, finite impulse response (FIR) filter to band-pass the EEG and sEMG signals between 1 and 100 Hz. We also applied FIR notch filters to reduce line noise at 50 Hz and the harmonic at 100 Hz. We re-referenced the EEG recordings to the average across all channels and removed electrooculography (EOG) contamination through the FastICA algorithm [36]. We then filtered the EEG and sEMG data into six bands: delta (1-4Hz), theta (4-8Hz), alpha (8-13Hz), beta (13-30Hz), low-gamma (30-50Hz), and high-gamma (50-100Hz).

In Experiment 1, we epoched the production period from -1 to 5s relative to the onset of the participant’s vocalisation. These onsets were detected from speech recordings for each trial as follows. We band-passed the vocal recordings between 95 and 300 Hz, a common range for human speech fundamental frequencies. We then computed an energy-based novelty function, as the first-order difference of the log-transformed audio signal, following convolution with a Hanning window (Section 6.1.1 in [37]). For each trial, we then defined the vocalisation production onset as the first local peak of the novelty function with a height greater than 1% of the maximum value over a period between -0.5 and 4 s from the production onset cue (Figure 1). In Experiment 2, we used only imagined vocalisation trials from the last five blocks. We epoched the recordings from -1 to 7s with regards to the preparation cue onset (Figure 1), the production period corresponded to 2 to 6 s of the epochs.

We removed the baselines (-1 to 0s) from all epochs and down-sampled the signals from 2048 to 256 samples per second. All trials in which the participants incorrectly recognised the emotion of the vocalisation were removed from further analysis. However, very few trials were removed; the overall error rate was 3.55%.

### 2.2 EEG-based emotion classification

#### 2.2.1 Feature extraction

We extracted time-frequency features from EEG recordings. Subsequent to epoching, we applied channel-wise Hilbert transforms to compute instantaneous band power on each of the six frequency bands. We first obtained the complex-valued analytic signal from the transform. The envelope of the EEG signals were then estimated by taking the root sum of squares of the real and imaginary parts of the analytic signal at each timestamp (equation 18.76 in [38]). We took a natural log transform on the square of the envelope, as an estimate of the EEG signal’s log-scaled instantaneous power for the corresponding frequency bands. The Hilbert transforms were done using an open-sourced Python implementation in scipy [39]. Finally, we computed the mean log-scaled power for each frequency band and each channel over 500 ms windows with 50% overlaps throughout the production period. This resulted in a time-frequency feature matrix of 6 frequency bands ×64 channels × 15 windows for each trial (0-500 ms, 250-750 ms, …, 3500-4000 ms relative to the production period onset). The 500-ms window with 50% overlap was chosen to capture temporal dynamics while keeping the feature matrix compact to reduce overfitting risk. This configuration aligns with prior EEG emotion-decoding studies and is standard in band-power feature extraction [40, 41, 42, 43]. A sensitivity analysis showed that classification accuracy remained stable across window sizes from 100 to 1000 ms (Appendix A.3).

#### 2.2.2 Cross-validation and chance-level accuracy

To verify whether emotion could be decoded from overt and imagined vocalisation EEG recordings, we performed within-participant single-trial classification of emotion categories for Experiments 1 and 2, separately. We set up a 25-fold cross-validation (CV) framework. In each fold, we picked one male and one female from the ten actors employed in the MAV dataset and selected trials in which presented vocalisations from the two speakers as the testing set. The remaining trials were placed in the training set. We then trained a logistic regression classifier using the training set and decoded the trials in the testing set. The default regularisation parameter C=1 (scikit-learn [44]) was used as it yielded stable performances (see Appendix A.2 for a complete analysis of the effect of C on classification accuracies). We standardised each feature using training-set statistics (mean and standard deviation), applying the resulting transformation to both the training and testing sets before classification. We then computed the classification accuracy as the proportion of correctly decoded trials over all trials in the testing set. We repeated the same process for all possible 25 combinations of male and female speakers. Subsequently, we constructed the CV framework and computed the accuracies on all participants.

We performed a shuffle test, repeated 100 times, to obtain an estimated distribution of chance-level classification accuracies. In each repeat, we randomly shuffled the training labels in all CV folds, ran the classifications, and obtained one fold-averaged chance-level classification accuracy. The classification accuracy observed on the original data was said to be statistically significantly greater than chance if it was greater than or equal to 95% of the accuracies in the corresponding distribution (p ≤0.05). The p-value was computed as *p* = (*N*_*H*_ +1)*/*(*N*_*T*_ +1), where *N*_*H*_ denotes the number of repeats where the accuracy was greater or equal to the observed accuracy on the original data, and *N*_*T*_ denotes the total number of repeats.

#### 2.2.3 Frequency band- and channel-wise feature importance

To examine the important frequency bands and EEG channels for emotion decoding, we performed frequency band-wise and channel-wise classification. We first computed the time-frequency features for all time intervals (“windows”), frequency bands, and channels (Section 2.2.1), yielding a flattened feature vector of length 5760 (15 windows ×6 frequency bands ×64 channels) for each trial. After evaluating classification accuracies on this full feature set, we evaluated classifiers on individual frequency bands (corresponding to feature vectors of length 960, that is, 15 windows ×64 channels). The frequency band with the highest accuracy underwent a channel-wise classification: we retrieved feature vectors belonging to one channel, evaluated the classification accuracy, and repeated the same process on the remaining 63 channels. Furthermore, we also computed F1-scores for each emotion category, to reveal the important channels specific to each emotion category. Specifically, for a given emotion category, we computed the number of true positives (TP), false positives (FP) and false negatives (FN) for the classifier’s prediction on trials belonging to this emotion, and computed the F1-score as 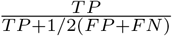 We repeated the same process on all CV folds and computed the fold-averaged F1-scores. We averaged the accuracy scores and F1-scores across participants. These measurements indicated the importance of frequency bands and EEG channels.

### 2.3 Examining EMG contamination

#### 2.3.1 EEG-sEMG channel-wise instantaneous correlation

We estimated the extent of EMG contamination in each EEG channel by examining synchronisation between EEG and sEMG recordings. EMG contaminates EEG primarily through volume conduction – the instantaneous spread of electrical activity through surrounding tissues [45, 46]. Therefore, we chose to quantify EMG contamination using absolute zero-lag Spearman’s *ρ*, as this measure is known to be highly sensitive to volume conduction [45]. Specifically, for each participant, we computed the absolute Spearman’s *ρ* between the instantaneous log-scaled high-gamma band power on each pair of sEMG and EEG channels after concatenating all trials (high-gamma band is the band with the highest classification accuracy, see Section 2.2.3 and 3.1). We then took a cross-participant average; a larger value indicated a larger EMG contamination in the corresponding EEG channel from the muscle groups covered by the corresponding sEMG channel. For each EEG channel, we also took the average of the absolute *ρ* statistics to the four sEMG channels to indicate an overall contamination level for the EEG channel. To establish a baseline for comparison, we shuffled the sEMG instantaneous band power across timestamps and trials, within channels and participants. We computed correlation coefficients between the sEMG and EEG channels after shuffling as described above, repeated the process 100 times, and took the average across repeats as the baseline.

#### 2.3.2 EEG versus sEMG versus combined features for classification

Further corroborative evidence for EMG’s contamination of EEG recordings is provided by a comparison of the emotion classification accuracy using sEMG features only versus combined EEG and sEMG features. We reasoned that, if EEG is free of EMG contamination and captures only cortical activity, then combining EEG and sEMG features should result in a higher classification accuracy. With high levels of EMG contamination, we expect the classification accuracies (EEG only versus combined EEG and sEMG) to be similar. We performed the above-described analysis using EEG and sEMG data recorded during the imagined production period on five participants in Experiment 2. We extracted sEMG features in the same way as that for EEG (see Section 2.2.1). Then, we compared classification accuracies on sEMG features only versus combined EEG and sEMG features. Additionally, we performed classification on EEG features only for comparison. Classification was performed using features from all frequency bands, or using only high-gamma-band features (that is, the band with the highest classification accuracy; see Section 2.2.3 and 3.1). The Nadeau & Bengio corrected t-test with significance level = 10% [47] and a paired Student’s two-tailed t-test were used to compare the classification methods within- and across-participants, respectively. We applied the Benjamini-Hochberg procedure to control the false discovery rate in these statistical tests [48].

### 2.4 Railroad cross-tie pattern

The railroad cross-tie pattern is a form of EMG contamination in scalp EEG, first described decades ago [49, 7] but still understudied. This pattern is of particular concern because it can spread across large areas of the scalp and can appear even during muscle relaxation. The railroad cross-tie pattern is characterised by fast, strong “spikes”, occurring at a rate of 8-18 “spikes” per second, at a significantly larger amplitude than spontaneous EEG activity [7]. Goncharova et al. showed that this pattern had a lateral distribution on the scalp over the temporal regions, and reported the occurrance of this pattern in EEG recordings not only when nearby muscles contracted, but also during relaxation (i.e., “resting-state”) [7]. We hypothesised that this pattern would also be present during imagined vocalisation. Here, we performed systematic detection of the railroad cross-tie pattern and analysed its differences between emotion categories.

#### 2.4.1 Pattern detection

The railroad cross-tie pattern stands out from high-gamma band EEG recordings in a way that is similar to that of neuronal action potentials in intracortical recordings. Therefore, we adopted a method commonly used to detect neuronal spikes in intracortical recordings [50, 51]. Specifically, from the imagined vocalisation production epochs (2 to 6 s after preparation cue onsets) in Experiments 2, we computed a threshold, *θ*, for each participant and each channel, using EEG recordings band-pass filtered between 50 and 100 Hz:

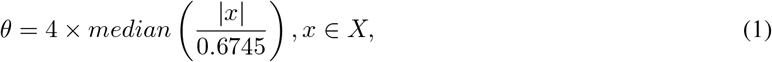

where *x* denotes an individual EEG sample and *X* denotes the set of all samples in the band-pass-filtered EEG recording for a given participant and channel.

We then detected all “spikes” with an absolute amplitude exceeding this threshold. We applied a refractory period of 50 ms (13 samples), thus allowing a maximum of 20 “spikes” to be detected per second. This process was repeated for all channels and participants.

#### 2.4.2 Statistical analysis on railroad cross-tie pattern

We applied statistical analyses to determine whether there exists an effect of emotion on the railroad cross-tie pattern. We computed the number of “spikes” per trial (NoS/T) for each trial, indicating the strength of the railroad cross-tie pattern on one channel. We then computed the mean NoS/T for five emotion categories separately, for each channel and participant. Next, we applied a repeated-measures ANOVA on the mean NoS/T using emotion as the within-subject factor, for each channel separately. We applied Bonferroni correction to control the family-wise error rate (FWER) across multiple channels [52, 53]. Over the channels with null hypothesis rejected under the ANOVA analysis, we performed a post-hoc analysis on the mean NoS/T values averaged across channels, using paired t-tests. The false discovery rate was controlled using the Benjamini-Hochberg procedure [48].

## 3 Results

### 3.1 Classification accuracies and important frequency bands/channels

All participants capably performed the emotion recognition task, achieving low error rates. On average, each participant incorrectly recognised emotion on 9.2 of 300 trials (3.1%) in Experiment 1 (overt vocalisation), and 9.8 of 250 (200 for participant P4) trials (3.9%) in Experiment 2 (imagined vocalisation).

After removing the above-mentioned error trials, we performed single trial classification on EEG recordings during both overt and imagined vocalisation. We extracted frequency band power features in overlapping 500 ms windows and examined the classification accuracy on individual bands (i.e., delta, theta, alpha, beta, low-gamma and high-gamma) and on combined features over all bands. Classification results showed that emotion could be decoded from EEG recorded not only during overt vocalisation, but also imagined vocalisation (all frequency-band features; overt, 78.1% accuracy; imagined, 36.4%; chance = 20%). For overt vocalisations, all participants demonstrated statistically significant classification accuracies above chance, regardless of the frequency band used. For imagined vocalisations, classification accuracies statistically significantly outperformed chance on most participants on alpha (17 out of 21 participants), beta (15/21), low-gamma (17/21) and high-gamma (16/21) bands. Overall, classification accuracies increased from lower to higher frequency bands and peaked in the high-gamma band both for overt (79.5%) and for imagined (37.0%) vocalisations. Additional analyses showed that imagined vocalisation classification accuracy was not correlated with participant age, and including the two older participants did not affect the results (Appendix A.1). Figure 3 summarises the classification results.

**Figure 3.**
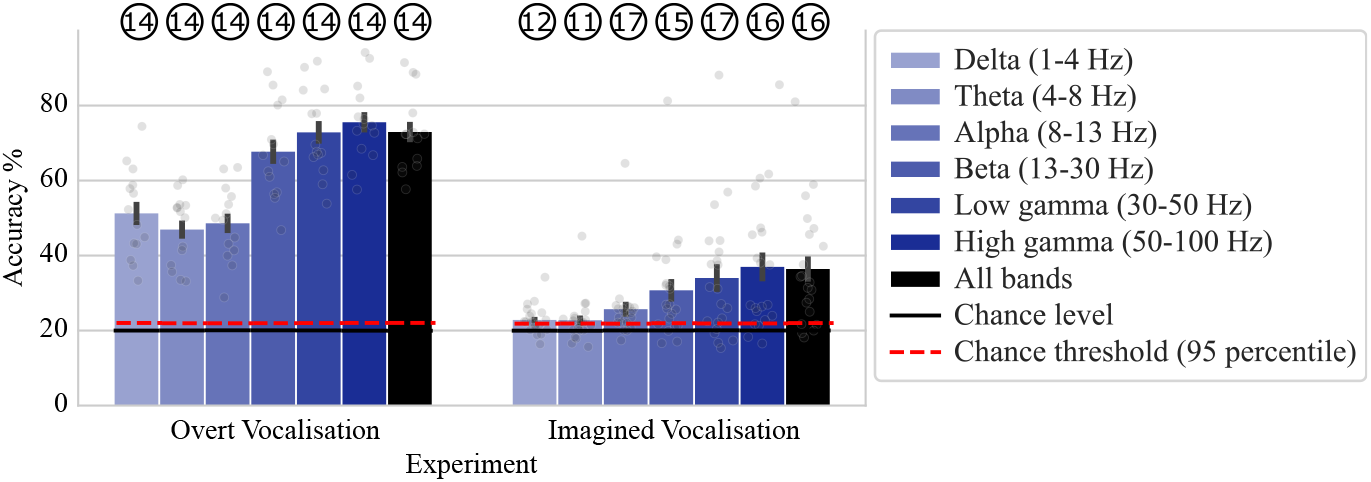
Emotion could be decoded from EEG recordings during both overt and imagined vocalisation; higher frequency bands resulted in higher decoding accuracies. The plot shows classification accuracies (%) for decoding emotion using log-scaled, windowed frequency band power features (Section 2.2.1). Error bars represent the standard error (SE) across participants. The numbers inside circles represent the number of participants (out of a total of 14 in overt and 21 in imagined vocalisation experiment) for whom the observed classification accuracy was statistically significantly greater than chance.

We performed channel-wise classifications on the most important frequency band – the high-gamma band. We computed classification accuracies and emotion category-wise F1-scores. We consistently observed higher classification accuracies on lateral EEG channels for both overt and imagined vocalisations. Specifically, EEG recordings for happy vocalisation trials exhibited significantly greater F1-scores in lateral channels over the temporal cortex (i.e., T7, T8 and nearby channels), highlighting their importance in emotion decoding. Figure 4 summarises the classification scores.

**Figure 4.**
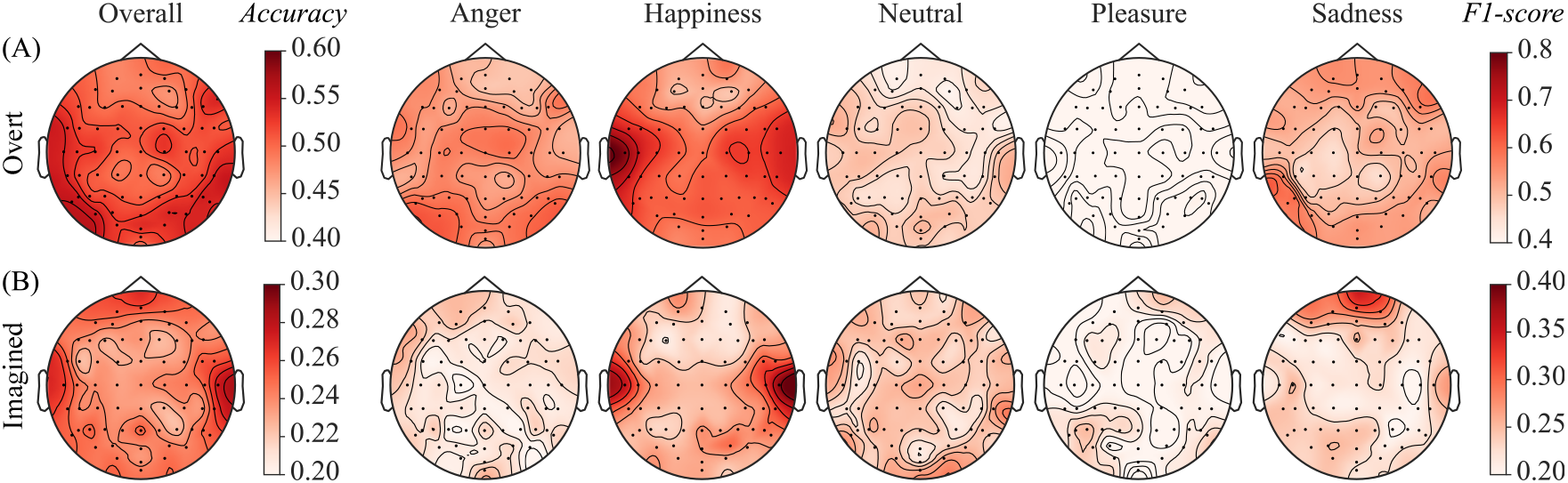
Channel-wise classification analyses using the high-gamma band showed similar channel importance patterns across A) overt and B) imagined vocalisation productions. The two left-most plots show overall, channel-wise classification accuracy averaged across participants; the ten other plots show, for each emotion, channel-wise F1-scores averaged across participants. On both overt and imagined vocalisations there was a prominent lateral pattern, especially for happy vocalisations.

### 3.2 EEG-sEMG correlations

For five participants in Experiment 2, we examined the instantaneous correlation between each EEG-sEMG channel pair. We computed log-scaled instantaneous high-gamma band power on EEG and sEMG channels and computed the absolute Spearman’s *ρ* between each pair of channels across trials (Section 2.3.1). We also obtained the baseline correlations in comparison by computing the statistics after shuffling samples across trials and time. Figure 5 shows the correlation statistics averaged across participants, as well as the shuffled baseline for comparison. We observed robust correlations between EEG and sEMG channels while the baseline showed no correlation between channels. These correlations corroborate EMG contamination in EEG and showed consistency with the spectral and spatial patterns observed across the full cohort (Section 3.1). The sEMG channels over the left medial frontalis region and the left temporalis region closely correlated with the nearby pre-frontal and lateral EEG channels as expected. Correlations were also observed between the left orbicularis oculi and zygomatic major sEMG channels and lateral EEG channels (especially T7 and T8). This pattern of correlation matched the pattern for channel-wise feature importance, especially in happiness emotion category (Figure 4).

**Figure 5.**
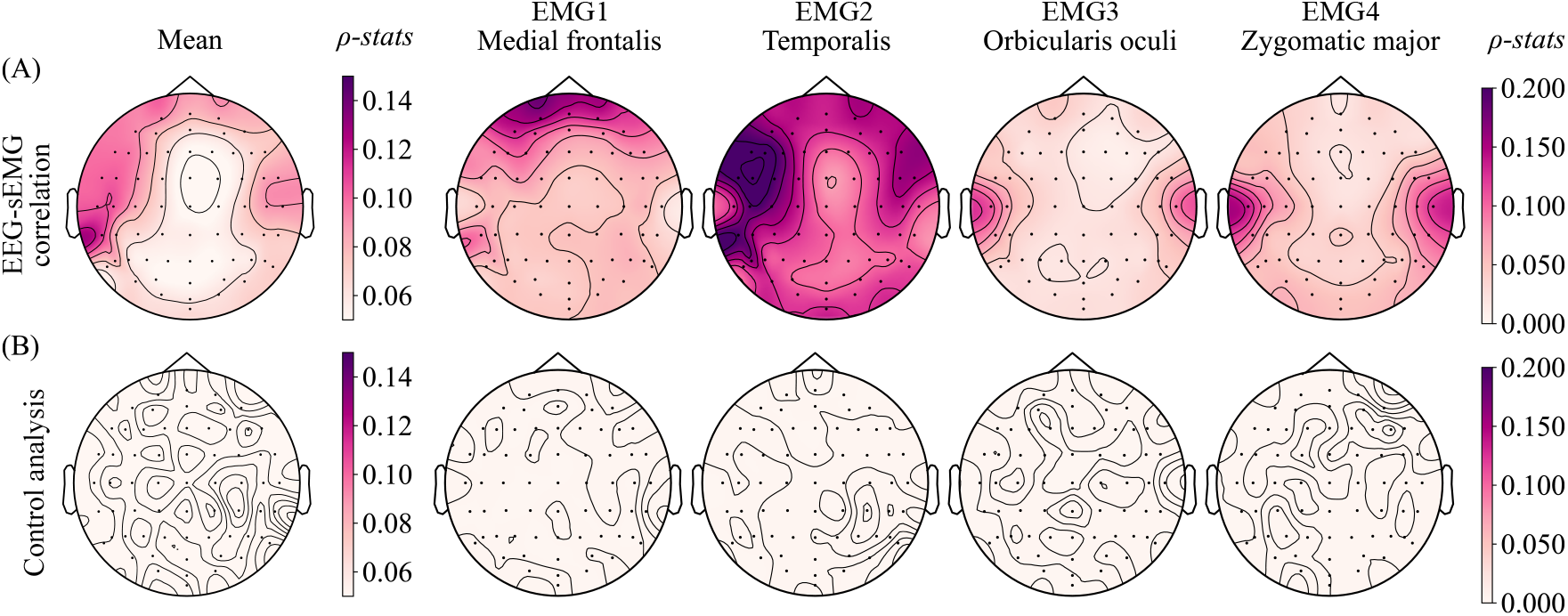
EEG activity was instantaneously correlated to the sEMG activity and showed distinct spatial patterns. We computed the absolute Spearman’s *ρ* correlation statistics for log-scaled instantaneous high-gamma band power between EEG channels and the four bipolar sEMG channels. Plots in (A) show the averaged statistics across participants and plots in (B) show the baseline statistics for comparison, obtained by shuffling EMG instantaneous power across trials and time. Notably, the correlations with the orbicularis oculi and zygomatic major regions were higher on lateral channels, matching that of the channel importance patterns for happy vocalisations (Figure 4).

### 3.3 EEG, sEMG and combined EEG-sEMG features decoding

We developed more evidence of EMG contamination by performing a comparison of classification accuracies using sEMG features only versus combined EEG and sEMG features. On five participants from Experiment 2 tasked with imagined emotional vocalisations, we performed classifications using EEG features alone, sEMG features alone, and using combined EEG and sEMG features. Overall, sEMG features alone yielded classification accuracies that were statistically significantly above chance. Adding EEG features, however, did not improve the classification accuracy. Within this subsample (n=5), this result suggests that EEG activity did not add extra information beyond sEMG activity, corroborating the existence of EMG contamination in EEG recordings. Using all frequency-band features, the accuracies did not significantly differ between feature sets across participants (34.9% for EEG, 39.3% for EMG, and 39.9% for combined features). However, within the high-gamma band, both sEMG and combined features achieved statistically significantly higher accuracies than EEG features alone (EMG v.s. EEG, 42.8% v.s. 34.5%, t=3.96, p=0.025, corrected; combined v.s. EEG, 43.5% v.s. 34.5%, t=4.04, p=0.025, corrected). Figure 6 summarises the classification accuracies.

**Figure 6.**
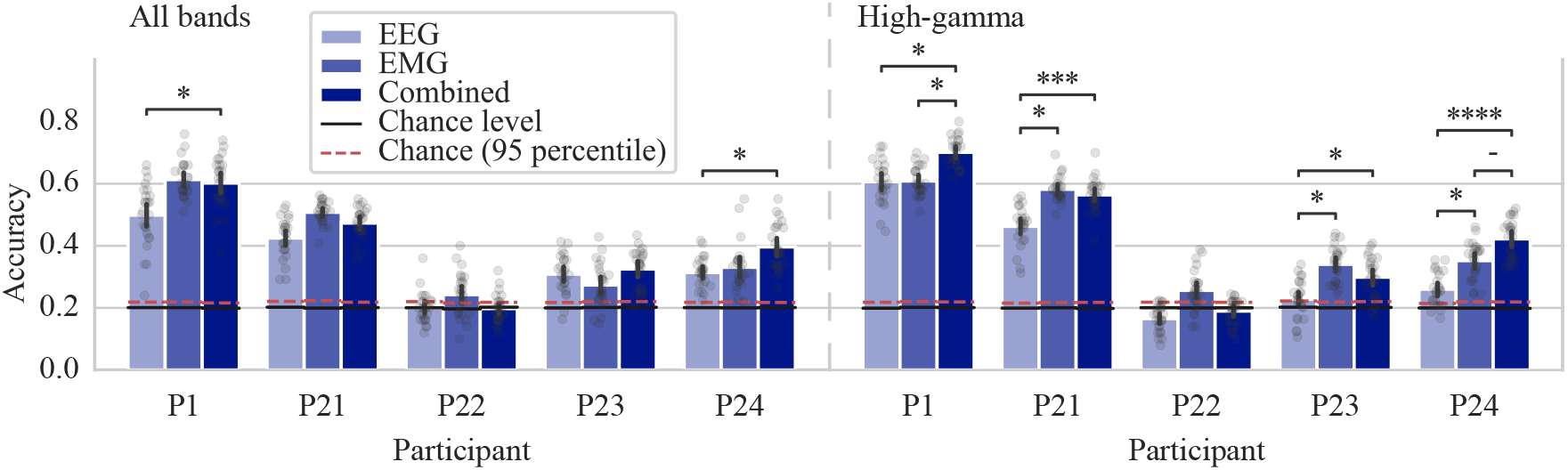
Overall, EEG features did not add extra information to sEMG features nor improved the classification accuracy on imagined vocalisations. The plot shows participant-wise classification accuracies using EEG, sEMG, and using combined EEG and sEMG features on all frequency bands and on high-gamma band only. Statistically significant p values after Benjamini-Hochberg correction are marked on the plot. Error bars show the 95% confidence intervals of the mean classification accuracies across 25 cross-validation folds. ‘-’ denotes 0.05*<*p*≤*0.1; ‘*’ denotes 0.01*<*p*≤*0.05; ‘**’ denotes 0.001*<*p*≤*0.01; ‘***’ denotes 0.0001*<*p*≤*0.001; ‘****’ denotes p*≤*0.0001.

### 3.4 Railroad cross-tie pattern

We observed the so-called railroad cross-tie pattern in EEG recordings during imagined emotional vocalisations [7]. In Figure 7, we illustrate two representative imagined vocalisation trials on a single channel EEG recording (i.e., FT7), one with and one without the railroad cross-tie pattern. The pattern, characterised by high-amplitude “spikes” occurring at a rate around 13 “spikes” per second in EEG recordings (band-passed between 50 and 100 Hz) and corresponding instantaneous high-gamma band power, was evident on channel FT7 on the sample happiness trial but absent in the sample neutral trial.

**Figure 7.**
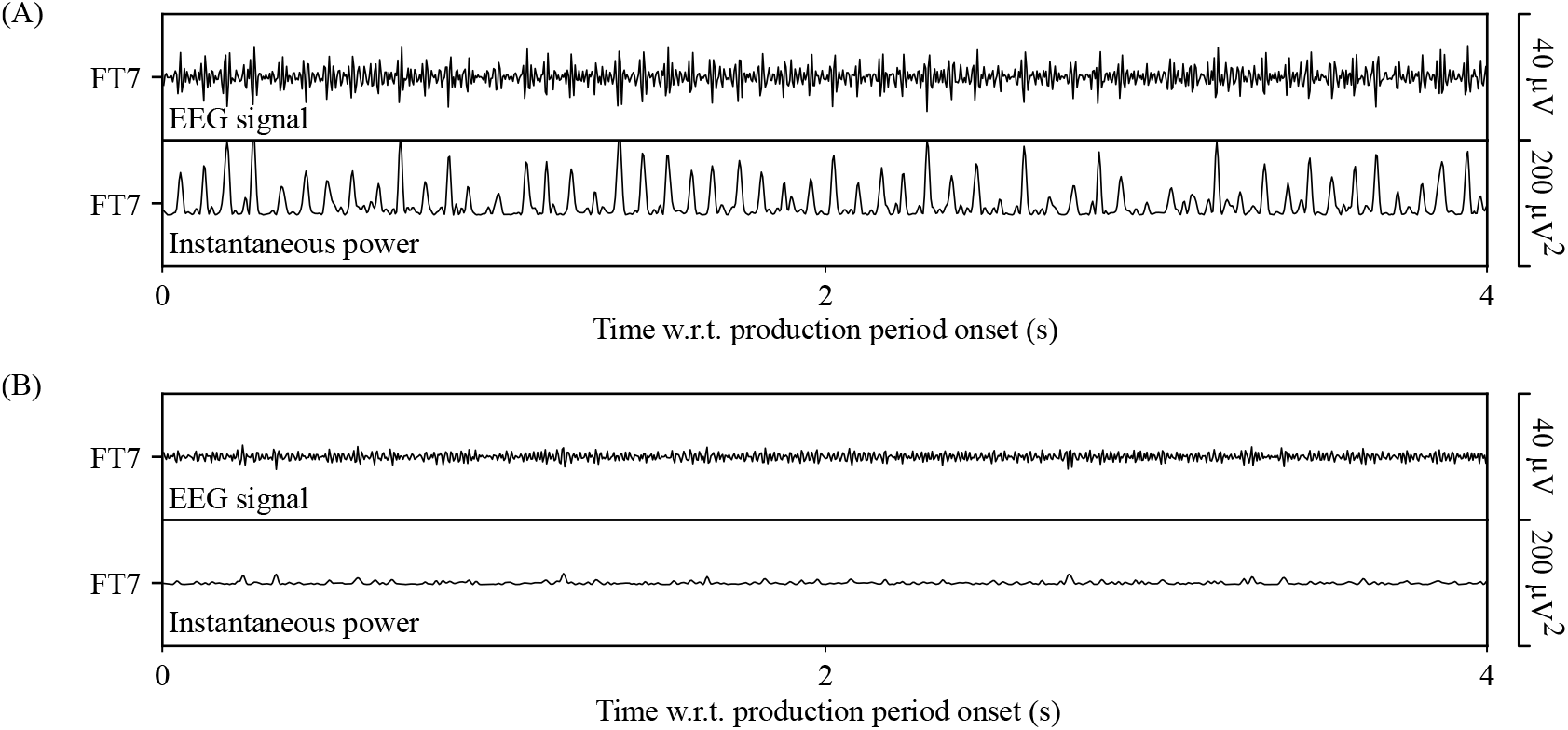
The railroad cross-tie pattern, a signature of EMG contamination, was present in EEG recordings, especially for “happiness” trials. The plots here show two representative trial EEG recordings (band-passed between 50 and 100 Hz) and instantaneous high-gamma band power from channel FT7 during imagined vocalisation production recorded on P17 in Experiment 2. Plot (A) shows one happiness trial with clearly visible railroad cross-tie pattern, characterised by sharp “spikes” in EEG recordings and in corresponding instantaneous high-gamma band power, occurring at a rate around 13 “spikes” per second, evenly spaced from each other. Plot (B) shows one neutral trial without the railroad cross-tie pattern.

We computed NoS/T as an indicator for the strength of the railroad cross-tie pattern. In twenty-one participants who performed imagined vocalisations, we then ran repeated-measures ANOVA and post-hoc tests to examine differences in NoS/T between emotion categories. The ANOVA analysis revealed emotion effects over five lateral channels including T7, F(4,80)=5.69, p=0.024, corrected; FT8, F(4,80)=6.22, p=0.013, corrected; FC6, F(4,80)=6.67, p=0.007, corrected; C6, F(4,80)=5.47, p=0.039, corrected; and T8, F(4,80)=7.12, p=0.004, corrected. Post-hoc analysis showed that over these channels, happiness trials had a larger mean NoS/T (15.56) than all other emotion categories (v.s. anger, 8.33, t=2.95, p=0.020, corrected; v.s. neutral, 8.01, t=2.95, p=0.020, corrected; v.s. pleasure, 9.46, t=3.79, p=0.011, corrected; v.s. sadness, 9.57, t=3.36, p=0.016, corrected). Results are presented in Figure 8. The greater NoS/T in happiness trial recordings resulted in greater high-gamma band power, and thus appeared to be used in classification to discriminate happiness trials from other. This inference is consistent with our observations that the lateral channels were more important for decoding emotion, particularly happiness trials (Figure 4). Additional analyses showed that NoS/T was not correlated with participant age, and including the two older participants did not affect the results (Appendix A.1).

**Figure 8.**
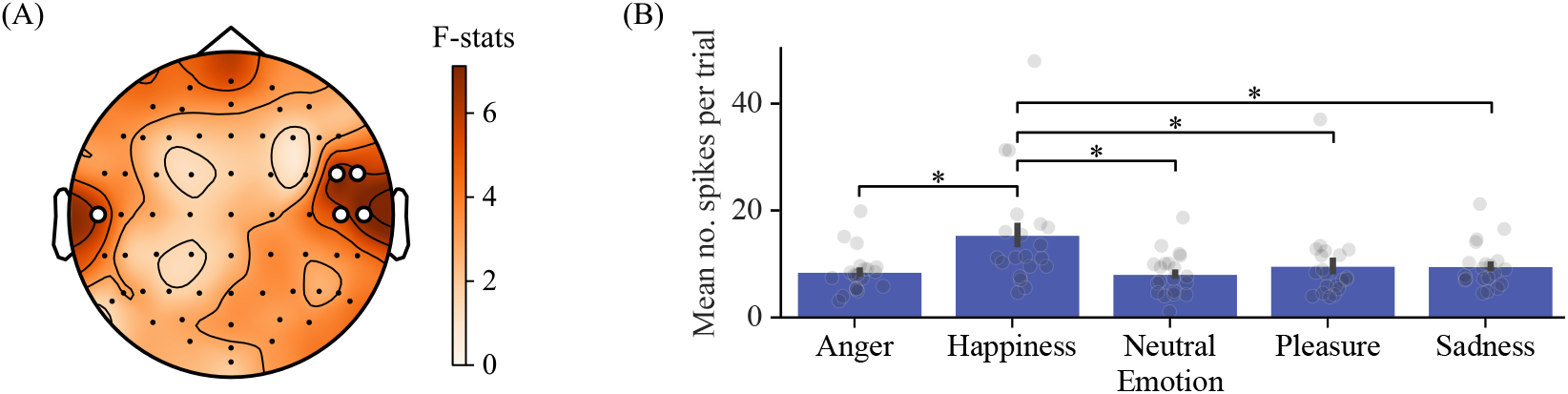
The strength of the railroad cross-tie patterns, computed using the number of (EMG-induced) “spikes” per trial (NoS/T), was greater in happiness trials than all other emotion categories. This difference was statistically significant over lateral channels. Plot (A) shows the repeated-measures ANOVA F-statistics on testing the emotion effect on mean NoS/T values across individual channels (statistically significant channels after a post-hoc Bonferroni correction are emphasised using white circles). Plot (B) shows the mean NoS/T values for each emotion category, averaged over the statistically significant channels. Error bars represent the standard errors (SE) across participants. ‘*’ denotes p*≤*0.05 for the post-hoc paired t-tests after a Benjamini-Hochberg correction.

## 4 Discussion

In this study, we decoded emotion from both overt and imagined vocalisations using instantaneous frequency-band power features extracted from EEG recordings. We determined the frequency bands and channels important for decoding. For both overt and imagined vocalisations, the higher frequency bands, especially high-gamma, provided the most important features; EEG channels over the temporal cortex (i.e., T7, T8 and nearby channels) were found to be most important for decoding emotion, particularly happiness. The consistency of the patterns between overt and imagined vocalisation across participants suggests a consistent source of decodable features. We hypothesised that this source was muscle activity, given its prominent presence during overt vocalisations. Moreover, the temporally distributed high-gamma activity closely matched a known muscle-induced EMG contamination pattern - railroad cross-tie pattern [7]. Analysis of simultaneously recorded sEMG supports our hypothesis. We observed an instantaneous correlation between sEMG and EEG activity in the high-gamma band over the lateral region of the scalp, matching feature importance patterns. When we combined EEG and sEMG features, classification accuracy was not improved when compared to sEMG features alone. We also identified the railroad cross-tie pattern in the EEG recordings during imagined vocalisations. This pattern was statistically significantly more prominent over the lateral channels, supporting classification of happiness trials.

Our results suggest that involuntary facial muscle contaminations during imagined emotional vocalisation production can be identified through EEG patterns, with corroborations from sEMG recordings. Specifically, high-frequency band power (high-gamma) over lateral channels near T7 and T8, modulated by increases in the railroad cross-tie pattern (NoS/T), should be considered as EMG contaminations rather than neural activity. A strong correlation in instantaneous high-gamma band power between facial sEMG and lateral EEG channels can corroborate that the observed band power feature originated from facial muscles and suggest responsible muscle groups. Furthermore, comparing classification performance using sEMG features alone versus combined EEG–sEMG features can provide additional corroboration. A similar accuracy to that obtained from EEG features alone supports the presence of EMG contamination.

Existing sEMG studies have reported involuntary activity of the zygomatic major and orbicularis oculi facial muscles during the perception of positive-valence stimuli. Increased activity was observed in the zygomatic major when participants were exposed to affective pictures of positive-valence [54, 55] or instructed to imagine a happy situation [56]. Cacioppo et al. also reported increased activity in the orbicularis oculi in response to affective pictures of positive-valence [54]. Our results confirm and extend these observations, suggesting that, during imagined happy vocalisations, the zygomatic major and obicularis oculi are involuntarily active, driving EMG activity – the railroad cross-tie pattern – over the lateral EEG channels.

While our results indicate that, for involuntary happy vocalisations, it is the railroad cross-tie pattern that contaminates EEG and drives classification, patterns for other emotions were not as clear. Nonetheless, our combined EEG and sEMG classification results suggest that these other emotions are likely associated with involuntary facial muscle activity which induces EMG patterns. For example, we observed greater importance of frontal channels (Fp1, Fp2 and nearby channels) for decoding sadness emotion (Figure 4). Multiple muscle groups near this region, including the frontalis (Figure 2) and the corrugator supercilii (muscle that pulls the brows together), have previously been found to be activated by stimuli of negative-valence, such as sadness [54, 57, 58, 59]. These involuntary muscle activations, often visually imperceptible, would generate EMG contamination on frontal EEG channels and could plausibly explain the channel importance maps we observed for sadness in this study.

Our finding that EMG’s contamination of EEG drives emotion decoding from scalp recordings has implications beyond imagined emotional vocalisations. The same spectral and spatial feature importance patterns, which here we attribute to muscle activity, have been commonly observed in EEG-based emotion decoding studies. The widely used SEED dataset is comprised of EEG recordings from participants tasked with watching emotional film clips [60]. Zheng and Lu decoded emotion from time-frequency EEG features and found that the beta and gamma bands and the lateral channels (T7, T8 and nearby channels) contributed the most discriminative features [60]. This pattern has been reported by many studies using the SEED dataset [61, 62, 63, 64, 65]. DEAP is another widely used dataset; this dataset is comprised of EEG recordings from participants tasked with watching emotional music videos [66]. Classification analyses of this dataset, too, commonly report the importance of high-frequency bands [66, 67, 68, 69]. Moreover, Yang et al. also reported the importance of high-frequency bands in their independently recorded dataset, which is comprised of EEG recordings from participants tasked with viewing affective pictures [70]. In these studies, the plausible presentation of EMG activity in EEG recordings was rarely mentioned. Low-pass filters were applied in all datasets, with cut-off frequencies at 50 Hz in SEED [60], 45 Hz in DEAP [66], and 80 Hz in Yang et al.’s study [70]. However, EMG and EEG spectra overlap at low frequencies [23, 24, 25, 26], and any unverified claim that low-pass filtering rejects EMG should be viewed with scepticism. Our results raise concern that involuntary facial muscle activity evoked by these emotional stimuli induced EMG contamination of EEG, and, in whole or in part, drove emotion classification. In other words, our results underscore the need for caution when interpreting (putative) EEG features and their association with emotion; the attribution of EEG features to the neural correlates of emotion is far from straightforward. Where feasible, sEMG should be recorded simultaneously with EEG to confirm the absence of muscle artefacts before making claims that EEG features reflect neural correlates of emotion. Notably, although sEMG was not recorded in SEED [60] or Yang et al.’s study [70], it was recorded over the zygomatic major and trapezius (neck) muscles in DEAP [66]; however, these sEMG recordings appear to have been rarely incorporated into analyses.

Recent works have also demonstrated that simultaneous sEMG recordings can inform and improve targeted EMG artefact removal [71, 72]. Beyond band-pass filtering, blind source separation (BSS) methods – most notably independent component analysis (ICA) and canonical correlation analysis (CCA) – have been developed to isolate artefacts, including facial muscle activity [73, 74, 75, 76]. ICA decomposes multi-channel EEG into statistically independent components with distinct topographies. Artefact-related components are typically identified manually, and the remaining components are then back-projected to reconstruct “cleaned” EEG [73, 74]. Manual identification is difficult and expert-dependent, while sEMG recordings have been shown to automate this process by forcing muscle activity into a small subset of components, thereby improving identification accuracy [71]. CCA decomposes EEG into maximally autocorrelated components and removes those with low autocorrelation, based on the premise that EMG is less auto-correlated than neural activity [75, 76]. However, the threshold for “low” autocorrelation is not clearly defined, and neural activity is often mixed with muscle activity in the same components and removed together [77, 72]. Mucarquer et al. [72] demonstrated that sEMG can be used as a “true” muscle reference to estimate and subtract muscle contributions within CCA components, thereby bypassing thresholding and better preserving neural signals.

Beyond artifact verification and removal, the use of EMG as a complementary or independent feature, rather than an artefact to be removed, may itself offer a practical benefit in BCI applications, as shown by our classification comparisons (Section 3.3). Recent studies have begun to combine EEG with other physiological signals in designing multi-modal human-computer interfaces, some of which combined EEG and EMG signals, and showed promising results in device control applications such as wheelchairs and exoskeletons [78, 79, 80, 81]. Since facial sEMG encodes emotion-dependent muscle activity, future BCI systems may benefit from including dedicated facial sEMG electrodes, which may yield a better signal-to-noise ratio than the residual EEG activity captured at standard scalp locations.

Our study is the first to demonstrate the modulation of involuntary muscle activity by emotion with identified EMG patterns during imagined emotional vocalisations — an important contribution to studies of emotional vocalisation and beyond, which nonetheless has limitations. One limitation of this study is that facial sEMG was recorded in only five of the 21 participants who performed imagined vocalisations. This small sample size constrains the statistical power of the EEG–sEMG correlation analysis and the combined EEG–sEMG classification comparison; although we think it likely, we have not demonstrated the generalisation of this result to all participants. Nevertheless, we note that our conclusions regarding EMG contamination do not rely solely on evidence from sEMG. Of the entire cohort of 21 participants, we observed that the spectral and spatial pattern of classification importance (high-gamma band, lateral channels) closely resembled the railroad cross-tie pattern. The characteristic high amplitude “spikes”, at a rate of around 13 per second, are consistent with motor unit firing rather than synchronous cortical oscillatory activity, supporting a myogenic rather than neural origin [7, 82, 83]. Furthermore, statistical analysis of spike counts across the full cohort (n=21) revealed a higher NoS/T during happiness trials. This increase in spike counts directly led to higher high gamma band power over the lateral channels, supporting our claim that such contamination contributes to emotion decoding performance. The sEMG findings, while limited in sample size, corroborated this interpretation and additionally suggested candidate source muscles (zygomatic major, orbicularis oculi). Given the limited sample size, this muscle-specific attribution was suggestive rather than conclusive. Dedicated studies with a broader facial sEMG coverage and larger cohorts are needed to confirm the specific muscular origins of the observed contamination.

Our study used single vowels (/ah/) from the MAV dataset. Single vowels minimise articulatory variability and eliminate semantic content, both of which could introduce confounds in our analyses. However, this choice also limits our ability to assess the extent of EMG contamination in EEG recordings during more natural, real-world speech. Future work may extend analyses to more complex utterances (e.g., words, sentences) to evaluate whether the observed EMG-contamination patterns generalise to more realistic articulatory scenarios. Our study also recorded sEMG only from the left side of the face. Studies using sEMG have shown facial symmetry during emotional expression [84], however, human ratings indicate that facial expressions are slightly more pronounced on the left side [85, 86]. For this reason, we used all available bipolar recording channels to measure sEMG over multiple candidate emotion-related muscle groups (medial frontalis, temporalis, orbicularis oculi, zygomaticus major; [34]) on the left hemiface, thereby maximising sensitivity to any emotional facial expression. Our use of this unilateral montage precludes the assessment of asymmetry in involuntary facial activity during imagined emotional vocalisations. Although we observed lateral EEG importance and instantaneous sEMG-EEG correlations over both hemispheres (T7/T8 and adjacent temporal channels), the relative contributions of ipsilateral versus contralateral muscles to these patterns remain unknown. Future studies could measure bilateral sEMG to clarify bilateral contributions and assess whether asymmetry exists and differs across emotion categories.

Moreover, our Experiment 1 cohort had a substantial gender imbalance (11 males, 3 females). Previous work has reported gender differences in emotional expression and associated physiological arousal patterns [87, 88], raising concern that this imbalance could bias our overt vocalisation classification results. To assess this, we compared classification accuracies between genders within the Experiment 1 cohort. No differences were observed across any frequency band (Student t-tests, all p*>*0.44; see Appendix A.4). However, the small number of female participants (n=3) limits statistical power, and the null findings should be interpreted with caution.

Our study was also limited to healthy participants. For people with facial paralysis (who might benefit from a BCI), the assumption that imagined emotional vocalisations elicit involuntary facial EMG does not necessarily hold. Most facial paralysis is due to peripheral facial nerve pathology [89], where the presence of EMG activity depends on the degree of denervation. In cases of complete denervation, voluntary motor unit action potentials (MUAPs) are absent due to loss of neural excitability (ch. 4 in [90]). Under such conditions, the EMG patterns observed in our healthy cohort would not be expected. However, it is common for residual innervation to persist, such that what appears to be a completely paralysed muscle may still exhibit low-amplitude MUAPs detectable with electrophysiological methods [91, 92, 93, 94]. This raises the possibility that some EMG activity could still be present in people with facial paralysis during imagined emotional vocalisation, although with reduced amplitude and potentially altered spatial or spectral characteristics. Whether such residual activity is useful for emotion decoding from EEG or facial sEMG recordings remains an open question.

## 5 Conclusion

Emotion can be decoded from EEG recordings during both overt and imagined emotional vocalisations, with the high-gamma band and lateral channels being the most important features for classification. However, analysis of simultaneously recorded facial sEMG indicates that this decoding is largely driven by involuntary facial muscle activity rather than neural correlates of emotion. In particular, for happiness, the railroad cross-tie EMG pattern was significantly more prominent over lateral EEG channels. These findings raised broader concerns about the probable interpretation of high-gamma, laterally-distributed EEG features as neural signatures in existing emotion decoding literature, and highlight the need for incorporating facial sEMG recording alongside scalp EEG.

## Notes

### Competing Interest Statement

The authors have declared no competing interest.

### Summary of Updates

Manuscript has been accepted by JNE: https://doi.org/10.1088/1741-2552/aeb073

