## Appendices for "Involuntary facial muscle activity during imagined vocalisation contaminates EEG and enables emotion decoding"

### Appendix

#### A.1 Effect of Age on Emotion Decoding

We examined whether the wide age range in Experiment 2 (19–73 years, mean 28.5 ± 12.82) influenced either the classification accuracy or the strength of the railroad cross-tie EMG pattern. The wide age range was due to two outlier elderly participants (59 and 73 years). The age range without these two outliers was 19–32 years (mean 24.5 ± 3.7).

Across all 21 participants in Experiment 2, we computed the Spearman correlation between age and high-gamma band classification accuracy during imagined vocalisation production. The correlation was not statistically significant (Spearman $\rho = -0.09$, $p = 0.698$), indicating that age did not systematically modulate emotion decoding accuracy (Figure A1). Removing the two oldest participants (ages 59 and 73) resulted in a comparable classification accuracy (37.0% before versus 37.3% after removal).

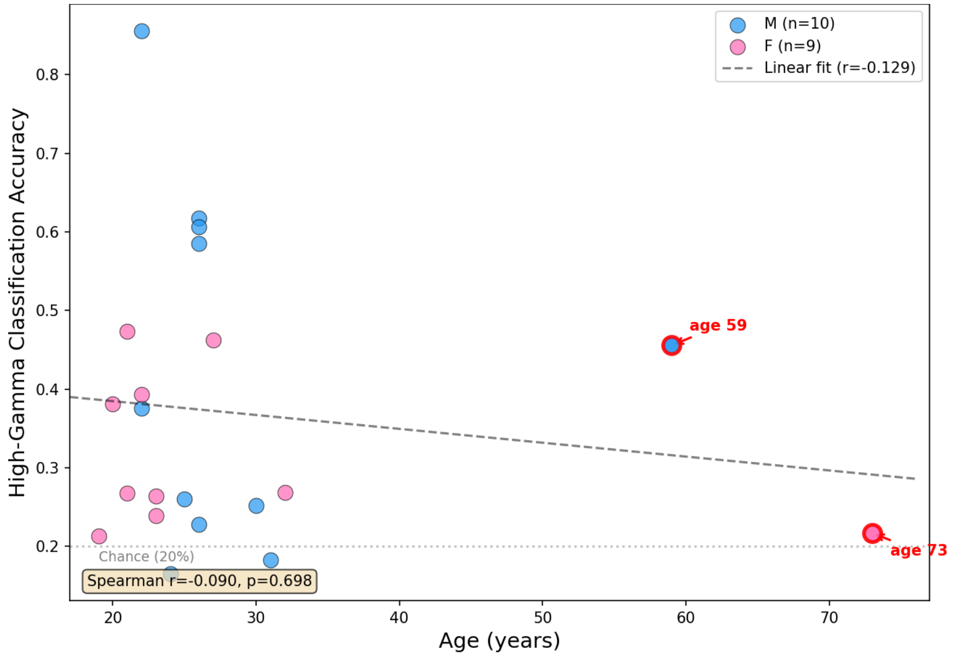


**Figure A1.** Age versus high-gamma band classification accuracy for imagined vocalisation (Experiment 2, N = 21). Blue markers indicate male participants; pink markers indicate female participants. The dashed line shows the linear fit. The grey dotted line indicates the classification chance level (20%).

We additionally examined whether age correlated with the strength of the railroad cross-tie pattern, quantified as the number of spikes per trial (NoS/T). For each participant, we used the mean NoS/T averaged across all five emotion categories and across the five lateral channels identified as significant in the main analysis (T7, FT8, FC6, C6, T8). The correlation between age and NoS/T was not statistically significant ($\rho = 0.174$, $p = 0.462$). This null result was confirmed across all five lateral channels individually (T7: $\rho = -0.263$, $p = 0.262$; FT8: $\rho = 0.296$, $p = 0.205$; FC6: $\rho = 0.329$, $p = 0.157$; C6: $\rho = 0.290$, $p = 0.215$; T8: $\rho = -0.060$, $p = 0.800$) and across all five emotion categories separately (all $p > 0.124$). Removing the two oldest participants as a sensitivity check did not change the correlation ($\rho = 0.189$, $p = 0.453$).

To verify that the emotion-specific railroad cross-tie pattern differences reported in the main analysis are not driven by the two oldest participants, we recomputed the post-hoc paired t-tests on the five significant lateral channels (T7, FT8, FC6, C6, T8) after excluding the two outliers. The results remained comparable. Across all five channels, happiness trials showed significantly greater NoS/T than all other emotion categories (all $p < 0.05$; Table A1), consistent with the main analysis. Together, these results indicate that age did not systematically modulate either the classification accuracy or the railroad cross-tie EMG contamination pattern.

**Table A1.** Post-hoc paired t-tests comparing the number of spikes per trial (NoS/T) for happiness versus each other emotion category on the five significant lateral channels, excluding the two oldest participants (N = 19). All comparisons remain statistically significant ($p < 0.05$).

| Channel | vs. anger | vs. neutral | vs. pleasure | vs. sadness |
|---|---|---|---|---|
| T7 | t=2.40, p=0.028 | t=2.59, p=0.018 | t=3.05, p=0.007 | t=2.51, p=0.022 |
| FT8 | t=2.40, p=0.027 | t=2.43, p=0.026 | t=2.93, p=0.009 | t=2.54, p=0.020 |
| FC6 | t=2.71, p=0.015 | t=2.44, p=0.025 | t=3.41, p=0.003 | t=2.65, p=0.016 |
| C6 | t=2.35, p=0.030 | t=2.30, p=0.034 | t=3.93, p=0.001 | t=2.70, p=0.015 |
| T8 | t=2.82, p=0.011 | t=2.72, p=0.014 | t=3.22, p=0.005 | t=3.44, p=0.003 |

#### A.2 Effect of Regularisation Parameter C

The logistic regression classifier used in the main analyses employed default regularisation (C = 1). To verify that results do not depend on this choice, we performed a full grid search over $C \in \{10^{-3}, 10^{-2}, 10^{-1}, 1, 10, 10^{2}, 10^{3}\}$ for each frequency band using the same 25-fold cross-validation framework (Section 2.2.2) on all 21 participants in Experiment 2. The accuracy remained stable across different C values (difference < 1.6% within any frequency band). We used the default C = 1 in the main analysis. Mean classification accuracies and standard errors (SE) are shown in Table A2 and Figure A2.

**Table A2.** Classification accuracy (%; mean ± standard error, SE) at each regularisation parameter C, for all frequency bands (N = 21). Values are averaged across all participants. Accuracy remained stable across the range of C values tested, with a maximum difference of less than 1.6% within any band.

| **Band** | **$10^{-3}$** | **$10^{-2}$** | **$10^{-1}$** | **1** | **10** | **$10^{2}$** | **$10^{3}$** |
|---|---|---|---|---|---|---|---|
| Delta | 24.1 ± 1.1 | 23.6 ± 0.9 | 22.9 ± 0.8 | 22.8 ± 0.8 | 22.7 ± 0.8 | 22.6 ± 0.8 | 22.5 ± 0.8 |
| Theta | 23.7 ± 1.3 | 23.4 ± 1.3 | 23.0 ± 1.3 | 22.7 ± 1.3 | 22.6 ± 1.3 | 22.5 ± 1.3 | 22.4 ± 1.3 |
| Alpha | 26.4 ± 2.1 | 26.3 ± 2.1 | 25.9 ± 2.1 | 25.7 ± 2.0 | 25.6 ± 2.0 | 25.5 ± 2.0 | 25.5 ± 2.0 |
| Beta | 30.8 ± 2.9 | 31.2 ± 3.0 | 30.8 ± 3.1 | 30.7 ± 3.1 | 30.5 ± 3.1 | 30.5 ± 3.0 | 30.4 ± 3.0 |
| Low gamma | 33.6 ± 3.4 | 34.2 ± 3.7 | 34.1 ± 3.8 | 34.0 ± 3.8 | 34.0 ± 3.7 | 34.0 ± 3.7 | 33.9 ± 3.7 |
| High gamma | 35.3 ± 3.5 | 37.6 ± 3.8 | 37.6 ± 3.9 | 37.0 ± 3.9 | 36.9 ± 3.9 | 36.7 ± 3.9 | 36.7 ± 3.9 |

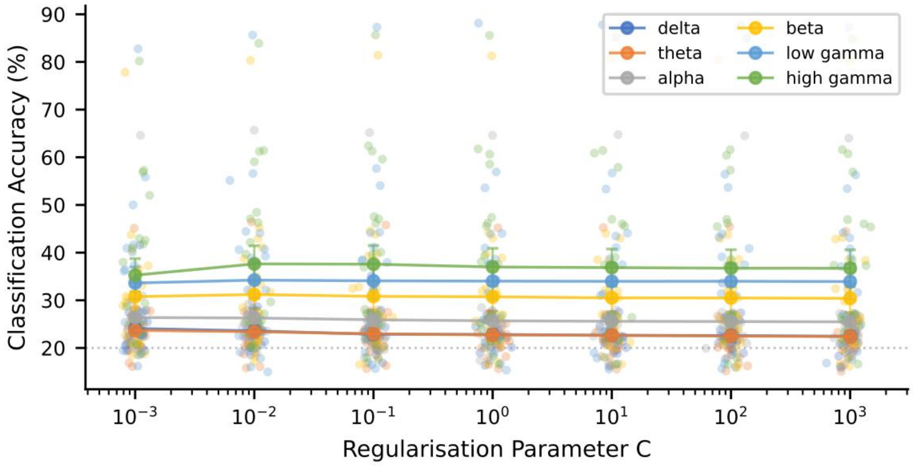


**Figure A2.** Classification accuracy (mean ± SE) across frequency bands at each regularisation parameter C. Lines connect mean accuracy across C values. Accuracy remained stable across $C \in \{10^{-3}, 10^{-2}, 10^{-1}, 1, 10, 10^{2}, 10^{3}\}$, with a maximum difference of less than 1.6% within any frequency band. The default C = 1 was used in the main analysis.

#### A.3 Effect of Time Window Size

We assessed the sensitivity of classification accuracy to the choice of EEG analysis window size. We extracted log-transformed high-gamma band power from EEG recordings and performed emotion decoding under the 25-fold cross-validation framework (Section 2.2.2) using five different window configurations: 100 ms (no overlap), and 250, 500, 750, and 1000 ms (all with 50% overlap). Accuracy was stable across all tested window sizes with a marginal increase at longer window sizes (Table A3 and Figure A3).

**Table A3.** Emotion classification accuracy (%; mean ± SE) by frequency band at five window sizes, for all 21 participants in Experiment 2. Accuracy was stable across all window sizes for all frequency bands, with a marginal increase at longer window sizes.

| Band | 100 ms | 250 ms | 500 ms | 750 ms | 1000 ms |
|---|---|---|---|---|---|
| Delta | 22.4 ± 0.9 | 22.7 ± 0.9 | 22.8 ± 0.8 | 22.8 ± 0.8 | 22.9 ± 0.9 |
| Theta | 22.3 ± 1.0 | 22.3 ± 1.2 | 22.7 ± 1.3 | 23.1 ± 1.3 | 22.9 ± 1.3 |
| Alpha | 24.7 ± 1.9 | 25.0 ± 2.1 | 25.7 ± 2.0 | 26.0 ± 2.1 | 26.1 ± 2.1 |
| Beta | 29.5 ± 2.6 | 30.6 ± 2.8 | 30.7 ± 3.1 | 31.2 ± 3.1 | 30.9 ± 3.0 |
| Low gamma | 33.2 ± 3.5 | 34.0 ± 3.7 | 34.0 ± 3.8 | 34.3 ± 3.7 | 34.3 ± 3.7 |
| High gamma | 35.8 ± 3.8 | 36.4 ± 3.8 | 37.0 ± 3.9 | 37.3 ± 4.0 | 38.0 ± 4.0 |

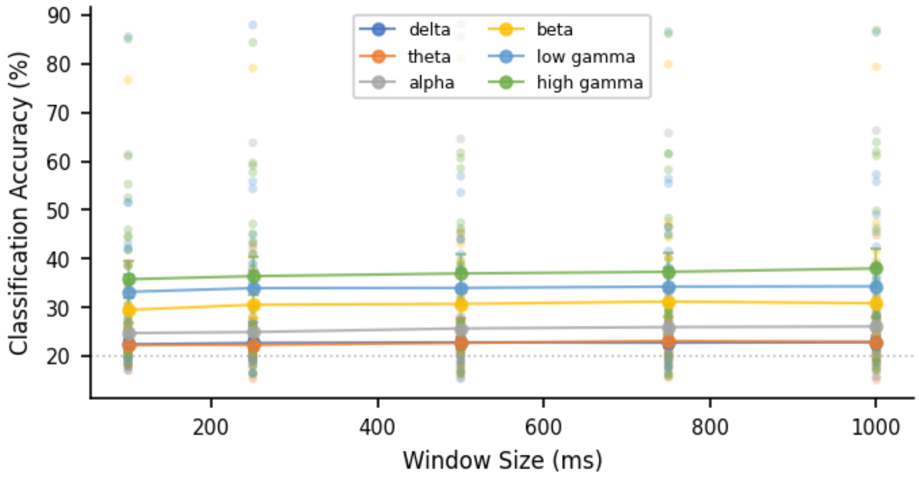


**Figure A3.** Emotion classification accuracy across frequency bands at five window sizes. The x-axis shows window size in milliseconds (100 ms, no overlap; 250, 500, 750, and 1000 ms, 50% overlap). Lines connect mean accuracy at each window size. Error bars represent ±1 SE. Accuracy was stable across all tested window sizes and all frequency bands, with a marginal increase at longer window sizes

#### A.4 Effect of Participant Gender

Experiment 1 (overt vocalisation, N = 14) had a severe gender imbalance (11 males and 3 females), potentially introducing bias in our classification analyses. We thus compared classification accuracies between male and female participants at all frequency bands. Student t-tests were performed on the classification accuracies between gender groups, within each frequency band.

Classification accuracies were comparable between male and female participants. No statistically significant differences were found in any band (all $p > 0.44$). However, these results should be interpreted with caution given the small number of female participants (Table A4 and Figure A4).

**Table A4.** Comparison of emotion classification accuracies between male and female participants in Experiment 1 (overt vocalization, N = 14, Male = 11, Female = 3). Classification accuracy mean ± SE are reported. Student t-tests were performed between gender groups within each frequency band. No statistically significant differences were found in any band.

| **Band** | **Male (% ± SE)** | **Female (% ± SE)** | **p (t-test)** |
|---|---|---|---|
| All bands | 76.9 ± 3.0 | 82.3 ± 6.6 | 0.444 |
| Delta | 51.8 ± 3.5 | 55.8 ± 9.3 | 0.634 |
| Theta | 46.0 ± 3.3 | 46.2 ± 7.8 | 0.985 |
| Alpha | 45.1 ± 3.4 | 50.8 ± 6.7 | 0.453 |
| Beta | 68.3 ± 3.4 | 72.6 ± 9.7 | 0.622 |
| Low gamma | 75.2 ± 3.1 | 79.7 ± 8.1 | 0.553 |
| High gamma | 79.2 ± 2.8 | 80.6 ± 9.3 | 0.840 |

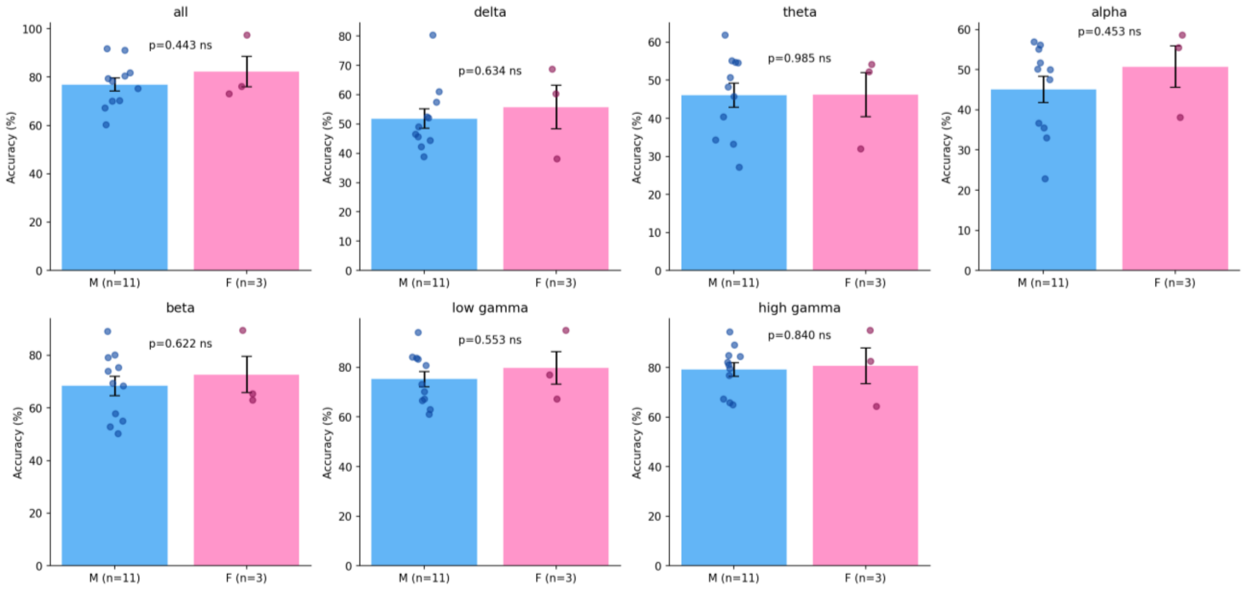


**Figure A4.** Experiment 1 (overt vocalisation) classification accuracy by gender for each frequency band. Individual participant data points are shown. Error bars represent ±1 SE. M: male (n = 11); F: female (n = 3). No statistically significant differences were found between gender groups in any frequency band.
